# Trophic flexibility favors coexistence of three aerial-hawking bat species in Venezuelan rice fields

**DOI:** 10.1101/2024.11.23.624971

**Authors:** Yara Azofeifa Romero, Jafet M Nassar, Jesús Mavárez

## Abstract

Coexistence among Neotropical insectivorous bats (IB) that share roosts, foraging areas, and prey likely relies on processes promoting trophic niche divergence. We examined the diet and activity times of three coexisting IB species (*Molossus molossus, Neoeptesicus furinalis*, and *Myotis nigricans*) in Venezuelan rice fields in Northwestern Llanos to assess dietary and temporal overlap. Using published sources, we categorized prey by hardness and flight speed, bats by bite force and functional group, and examined the relationships among these variables. IB species showed differences in prey composition, type, and activity timing. As expected from its fast flight and strong bite, *M. molossus’* diet consisted primarily of fast-flying, highly sclerotized insects, with activity times peaking significantly earlier than in the other two bat species. In contrast, *M. nigricans* and *N. furinalis* had diets consisting primarily of slower-flying prey and showed high temporal overlap in activity, although with different peak foraging times. Notably, *N. furinalis*’s stronger bite may have enabled it to exploit more sclerotized prey than *M. nigricans*, despite similar flight capabilities—indicating that prey hardness helps reduce dietary overlap. These findings suggest that fine-scale trophic niche partitioning, enhanced by the rich insect fauna of rice fields, facilitates the coexistence of these ecomorphologically distinct IB species.

Flexibilidad trófica favorece la coexistencia de tres especies de murciélagos cazadores aéreos en cultivos de arroz de Venezuela.

**Teaser Text:** We simultaneously identified prey items in feces and measured the activity times of three aerial-hawking bat species in Neotropical rice fields. We then explored the relationships between these variables and the functional traits of both prey and predators, confirming that coexistence might be facilitated by highly productive environments.

**RESUMEN:** La coexistencia entre murciélagos insectívoros (MI) neotropicales que comparten refugios, áreas de forrajeo y presas probablemente depende de procesos que promueven la divergencia del nicho trófico. Examinamos la dieta y los tiempos de actividad de tres especies coexistentes de MI (*Molossus molossus*, *Neoeptesicus furinalis* y *Myotis nigricans*) en arrozales de los Llanos Noroccidentales de Venezuela para evaluar superposición dietaria y temporal. Utilizando publicaciones, categorizamos a las presas según su dureza y velocidad de vuelo, a los murciélagos según su fuerza de mordida y grupo funcional, y examinamos las relaciones entre estas variables. Las especies de MI mostraron diferencias en la composición, tipo de presas y tiempo de actividad. Como se esperaba por su vuelo rápido y mordida fuerte, la dieta de *M. molossus* consistió principalmente en insectos de vuelo rápido y esclerotización alta, con tiempos de actividad que alcanzaron un máximo significativamente antes que en las otras dos especies de murciélagos. En contraste, *M. nigricans* and *N. furinalis* tuvieron dietas que consistieron principalmente en presas de vuelo lento que mostraron una alta superposición temporal en la actividad, aunque con distintos tiempos de forrajeo máximo. Cabe destacar que la mordida más fuerte de *N. furinalis* pudo haberle permitido explotar presas más esclerotizadas que *M. nigricans*, a pesar de sus capacidades de vuelo similares, lo que indica que la dureza de las presas ayuda a reducir la superposición dietaria. Estos hallazgos sugieren que una partición fina del nicho trófico, facilitada por la rica fauna de insectos en los cultivos de arroz, promueve la coexistencia de estas especies de MI ecomorfológicamente distintas.

**Palabras clave:** cultivos, humedales artificiales, metabarcoding, murciélagos insectívoros, tiempo de actividad.

## Background

Insectivorous bats (IB) are remarkably diverse predators with 20-45 species coexisting in syntopy in tropical forests and croplands in the Neotropics, Western Africa, and Southeast Asia (Kingston et al. 2003; Williams-Guillén and Perfecto 2011; Reardon and Schoeman 2017). The high local species richness of IB is likely a result of ecomorphological differences allowing for a wide variety of hunting and echolocation strategies among coexisting species, making possible trophic niche differentiation. Indeed, coexisting IB usually differ in wing morphology, body mass, skull shape, and echolocation call design, all of which correlate with habitat use, foraging strategies, and types of prey (Marinello and Bernard 2014). Accordingly, foraging site and diet represent important dimensions of ecological differentiation that contribute to resource partitioning among IB species.

Acoustic survey studies conducted in tropical and temperate environments have shown that sympatric aerial IB species partition foraging sites both horizontally and vertically, even at fine-grain spatial scales (Marques et al. 2016; Roemer et al. 2019; Beilke et al. 2021). In addition to spatial segregation, these species also partition resources along temporal dimensions, such as emergence time, timing of peak resource use (e.g. prey, water, or roost), and activity duration (Adams and Thibault 2006; Holland et al. 2011; Thomas and Jacobs 2013). However, how these temporal dimensions contribute to reducing interspecific competition is still unclear, particularly when less specialized or ecomorphologically distinct species coexist with dominant foragers (Mayberry et al. 2020; Beilke et al. 2021). Although the mechanisms behind spatiotemporal segregation remain understudied, such segregation promotes dietary divergence among IB species.

Clear differences in the diets of IB are not always evident, which is why increasingly precise dietary analyses are required to reveal underlying patterns of resource partitioning among coexisting species. Recent genetic metabarcoding analyses of diet in IB from temperate and tropical regions—conducted in diverse environments such as deserts, protected areas, agricultural landscapes, and settlements near caves— provide good evidence supporting trophic flexibility, with different degrees of dietary overlap that can be explained by subtle spatiotemporal partitioning of food resources (Bohmann et al. 2011; Salinas-Ramos et al. 2015; Gordon et al. 2019). On the other hand, larger dietary overlaps can be seen among IB species from highly productive periods or environments characterized by an elevated abundance of insects (Andriollo et al. 2021; Srilopan et al. 2024). Such conditions are typical of agricultural landscapes, where bats—capable of forming IB-rich ensembles—have become the focus of trophic niche partitioning studies that highlight their role as controllers of insect pests (Liu et al. 2024; Srilopan et al. 2024).

Rice fields provide an ideal setting for examining trophic niche partitioning, as their managed conditions create a humid, prey-rich environment that attracts ecomorphologically distinct IB species (Liu et al. 2024). Building on previous research in these rice fields (Azofeifa et al. 2019), this study provides unpublished dietary data on syntopic IB species to explore whether their close coexistence is facilitated by their differences in ecomorphological traits and activity times. With that aim, we performed DNA metabarcoding analysis of diets and assessed activity times inferred from echolocation calls, to examine trophic niche partitioning of three aerial-hawking IB species in rice fields in the Northwestern Llanos of Venezuela: *Molossus molossus* (Molossidae), *Myotis nigricans*, and *Neoeptesicus furinalis* (both Vespertilionidae).

These species are frequently sympatric, can inhabit the same roosts (Wilson and LaVal 1974) and forage in the same areas, including relatively open spaces, forest gaps and edges, and mosaics of agriculture and forestland in rural environments (Linares 1998). However, they differ in various morphological and behavioral traits that greatly affect their feeding strategies (Freeman 1981; Norberg and Rayner 1987). In this regard, *M. molossus* has a larger body, forearm, skull, and bite force than *M. nigricans*, while *N. furinalis* has values that are intermediate (e.g., body weight) or similar to *M. molossus* (e.g., forearm length and bite force) (Linares 1998; Aguirre et al. 2002). Additionally, differences in wing shape and aspect ratio imply a fast and high flight in uncluttered spaces in *M. molossus* (Marinello and Bernard 2014; Amador et al. 2020), but a comparatively slower and more maneuverable flight in background-cluttered spaces in *M. nigricans* and *N. furinalis* (Amador et al. 2020).

We predict that given the relatively larger size and distinctive flight type of *M. molossus*, its diet should be composed of a broader spectrum of prey and show relatively low overlap with the other two species. Under the same argument, the diets of *N. furinalis* and *M. nigricans* should reflect the differences in their body sizes, but also show some overlap given their similar flight types. Furthermore, the relatively strong skull, fast flight, and long-term calls of *M. molossus* should allow for a significant contribution of highly sclerotized and fast-flying insects in its diet. In contrast, these types of insects should have a lower frequency in the diets of *N. furinalis* and *M. nigricans*, being replaced by less sclerotized and slow-flying insects. Some predictions can also be made about activity times. For instance, assuming that activity times in these bats are maximal during periods of high availability of preferred prey, activity overlaps are expected to be high if prey flights are synchronous and low if they are not. According to this, *N. furinalis* and *M. nigricans* should exhibit significant activity overlap given their probably similar diets. For *M. molossus*, the overlap with the activity of other bats could be high if the maximum availability of preferred prey occurs at the same time for all three species, and relatively low if the preferred prey of *M. molossus* are asynchronous.

## MATERIALS AND METHODS

### Study area

This study was conducted in the Portuguesa state, part of the Northwestern Llanos of Venezuela, where two rice farms were selected: Colonia Agrícola Turén, Turén (hereafter “Turén” 9.215005267, −69.122405063, 140 m asl.) and Agropecuaria Durigua, Acarigua (hereafter “Acarigua” 9.542531825, −69.115494594, 180 m asl.) (Fig. 1). The two farms, located 45.6 km apart, lie in a flat, rice-dominated landscape with only 4–21% fragmented forest cover across 78.5 km² (Azofeifa et al. 2019).

**Fig. 1.**
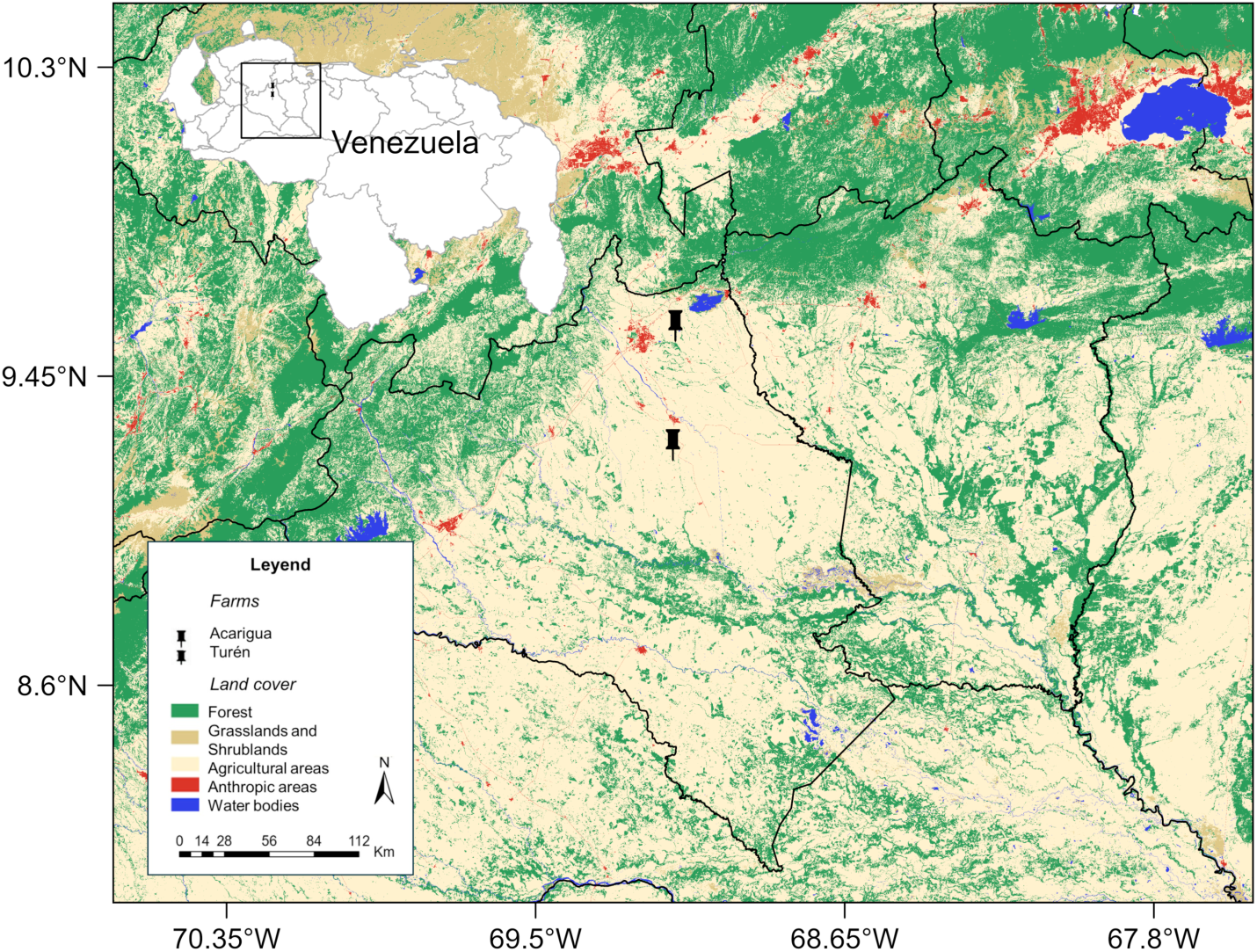
Map of Acarigua and Turén rice farms in the Northwestern Llanos of Portuguesa, Venezuela. The base map includes the 2014 land-cover layer from Red MapBiomas and Red Amazónica de Información Socioambiental Georreferenciada (RAISG).

### Sampling

#### Bats

Bat captures followed the guidelines recommended by the American Society of Mammalogists (Sikes et al. 2016). Between November 2013 and September 2014, each farm was visited six times, with ca. one-month interval between visits. Bats were captured using two 2.6 × 9.0 m mist nets (Avinet, Inc.), placed near roosting sites on two consecutive nights per visit at each site, and subsequently identified using the keys by Linares (1998) and Gardner (2007) (Supplementary Data SD1). In Turén, the three bat species cohabited beneath a human-made tent constructed with dried palm leaves, whereas in Acarigua, only *M. molossus* and *M. nigricans* were found roosting under the zinc roof and within the concrete beams of a warehouse. *Neoeptesicus furinalis* probably roosts in the forest remnants nearby this farm, as it was recorded foraging in the rice fields. Bats were captured between 20:00 and 21:30 h, shortly after the main peak of nightly foraging activity, as they returned to their roosts. Since the roosts were located in front of the rice fields and we consistently observed bats flying toward them, we assumed that their foraging activity was concentrated in these crops. The bats were placed inside a clean cotton bag to collect feces. Subsequently, fecal pellets collected were placed under a small cotton ball at the bottom of a sterile 15 mL polypropylene centrifuge tube, and dried by filling the tube with silica gel type III (Sigma-Aldrich H S7625-500 G). In total, fecal samples from 136 bats were collected: 71 *M. molossus*, 44 *M. nigricans*, 21 *N. furinalis*.

#### Insects

Nocturnal insects were captured using a white-light intersection trap placed on the rice fields on the same dates and hours of bat captures. The purpose of these collections was to create DNA reference libraries and to corroborate similarities among insect communities, as suggested by the literature on agricultural crops. Insects were identified by Moreno et al. (2020), preserved in 70% ethanol, and stored at room temperature until genetic analysis.

### DNA extractions

#### Bat fecal samples

DNA was extracted from the 10 largest fecal pellets of each bat sample using the DNasy Blood & Tissue Kit (Qiagen, Valencia, CA, USA). Sterile 1.5 mL microcentrifuge tubes containing fecal pellets and 1 mL lysis buffer solution were placed on a shaker at 30 rpm in an incubator at 55-56°C for two hours. Afterward, samples were centrifuged for 1 min at 1000 rpm, and 100 µL of supernatants were extracted and deposited in new sterile 2 mL microcentrifuge tubes. The remaining protocol steps followed the manufacturer’s instructions. To detect cross-contamination, extractions were performed in batches of 24 tubes, including two no-feces negative controls with and without water. Extracted DNA was stored at −18°C until subsequent PCR analysis.

#### Insects

DNA was extracted from two legs of insects with body size > 10 mm and from five whole bodies of smaller ones. Extractions were performed using the DNasy Blood & Tissue Kit following the manufacturer’s instructions.

### PCR amplification and sequencing

Two regions in the mitochondrial 16S rDNA unit were PCR-amplified for both fecal and insect samples: “Inse01” with primers Inse01-F 5’-RGACGAGAAGACCCTATARA-3’ and Inse01-R 5’-ACGCTGTTATCCCTAARGTA-3’, and “Arth02” with primers Arth02-F 5’-GATAGAAACCAACCTGGYT-3’ and LR-N-12945-2 5’-GCGACCTCGATGTTGGATT-3’ (Taberlet et al. 2018). All the primers had a common adapter used for Illumina sequencing and an individual 8-bp barcode that allowed the simultaneous sequencing of all samples.

DNA extracts were PCR-amplified in triplicate for fecal samples and once for insects. PCR reactions (volume 20 µL) contained: 10 µL of AmpliTaq Gold Master Mix (Applied Biosystems™, Foster City, USA), 0.2 µM of each primer, 5.84 µL of water, 0.04 µg of bovine serum albumin (Roche Diagnostic, Basel, Switzerland), and 2 µL of DNA diluted 1:20. The PCR protocols were: 1x: 10 min at 95 °C, 45x: 30 s at 95 °C, 30 s at 52°C (16S-1) or 50 °C (16S-2), 60 s at 72 °C, 1x: 7 min at 72 °C. The PCR batches included two extraction negatives, a PCR-negative (water), and a blank (empty well). The PCR products were pooled per replicate, resulting in eight pools: six bat diets (three per metabarcode) and two insects (one per metabarcode). Amplicon pools were purified using the MinElute PCR Purification kit (Qiagen), quantified using a Qubit 2.0 Fluorometer (Life Technology Corporation), and sent to Fasteris SA (Geneva, Switzerland) for library preparation and sequencing using the MetaFast protocol (www.fasteris.com/metafast; Taberlet et al. 2018).

### Bioinformatic analyses

The raw sequences were processed separately for each library using the *OBITools* package (Boyer et al. 2016). Reads were assembled with a minimum quality score of 40 and assigned to samples based on unique tag and primer combinations, allowing for two mismatches with the primer. Sequences more abundant in extraction, PCR and blank negative controls than in samples were considered as contaminants and removed. Sequences with coverage <10 reads, or with lengths outside the expected range (e.g., 76–168 bp for Arth02, Taberlet et al. 2018), or present in only one of the three fecal sample replicates were also discarded. The remaining sequences were clustered into prey MOTUs (*i.e*. Molecular Operational Taxonomic Unit) using the sumaclust algorithm with a sequence similarity of 97% (Mercier et al. 2013). Taxonomic assignment of MOTUs was achieved using the NCBI BLAST algorithm (Altschul et al. 1990), first against the 16S sequence database from local insects, and then against GenBank for the sequences that could not be assigned in the previous way. The following *ad hoc* assignment rules were applied in this study: (1) if there was a single best match: species level for similarity ≥ 98%, genus level for 95% ≤ similarity < 98%, family/sub-family level for 92% ≤ similarity < 95%, order level for similarity < 92%; (2) if there were multiple best matches: assigned to the lowest taxonomic level subsuming all best matches (e.g., assigned to the generic level if all best matches correspond to two or more species in the same genus). Sequences that remained unassigned were classified as “unknown”. A table of diet items was created by removing rare occurrences (frequency < 0.01%) and converting the counts into presence/absence values. We then calculated the percentage of prey by taxon (Pi) in the diet of three IB species as Pi = (∑Ni/N total) × 100, where ∑Ni is the total number of prey items for taxon i found in the diet and N total is the total number of prey items across all taxa in the diet.

### Bat activity time

We used two Echo Meter 3+ detectors placed on top of 2 m tall poles to record bat calls as described by Azofeifa et al. (2019). Bat calls were recorded continuously for 240 min, beginning at sunset. Each recording session occurred in the same two consecutive nights of bat capture mentioned above, resulting in four bat recordings per rice farm and session. This two-night recording scheme was therefore also repeated six times, with a one-month interval between two consecutive sampling sessions, resulting in 24 activity records per farm.

### Data analyses

Prior to dietary analyses, we used data from Moreno et al. (2020) to compare the insect communities between farms. We applied Hellinger transformations to the abundance data, calculated Bray-Curtis distances, and performed a PerMANOVA using PRIMER v7 software (Clarke and Gorley 2015). As insect communities did not differ between farms throughout the year (Pseudo-*F*_1,10_ = 1.234, *P*-perm = 0.234), neither within the dry (Pseudo-*F*_1,4_ = 1.033, *P*-perm = 0.598) nor the rainy seasons (Pseudo-*F*_1,4_ = 1.113, *P*-perm = 0.398), we combined the data from both farms for subsequent dietary analyses.

All diversity and composition analyses were based on presence/absence (incidence) data. Dietary diversity (Hill numbers q = 0, 1, 2) was quantified using iNEXT function in the R package *iNEXT* 3.0.1. Sample-size and coverage-based rarefaction/extrapolation curves were generated with ggiNEXT function. The percentage of sampling completeness for richness (q = 0) was calculated as (1-Q1/U) x 100, where Q1 is the number of prey singletons and U is the total number of prey observed (Hsieh et al. 2016). Differences in diet composition were analyzed with PerMANOVA tests based on Jaccard distance matrices using PRIMER v7 software (Clarke and Gorley 2015), followed by pairwise comparisons among bat species using non-parametric pseudo-*t* tests (akin to *t*-tests).

We employed Generalized Linear Models (GLMs) with a Poisson distribution, using the R package *MuMIn* 1.47.5 (Bartoń 2023), to assess the impact of three factors—IB species, functional group (uncluttered space *vs*. background-cluttered space), and bite force (high *vs*. low)— on prey characteristics such as the degree of sclerotization (strong, intermediate, or soft) and flight behavior (fast, intermediate, or slow) (Supplementary Data SD1 and SD2). The response variable in each model was the count of prey items within each category. We determined the relative importance of each predictor by comparing Akaike’s Information Criterion (AIC), calculating Akaike weights (wi), and examining the magnitude and direction of the estimated coefficients (β). Statistical significance was determined by *P* < 0.05, with confidence intervals excluding zero indicating a significant impact for the predictor.

Trophic niche breadth was estimated using Hill numbers of order q=1 and q=2 with the iNEXT function (Hsieh et al. 2016). Trophic niche overlap among the three bat species was assessed as pairwise Hill-number-based similarity (1-β_q_) for q=0, 1 and 2 using the hill_taxa_parti_pairwise function in the *hillR* package v. 0.5.3 (Li 2018).

Additionally, to visually illustrate dietary overlap, we conducted a PCA on the Hellinger-transformed incidence matrix using the rda function in the *vegan* package v. 2.6-10, which centers each prey variable to zero mean and applies a community-style PCA with equal Euclidean weighting (Oksanen et al. 2025).

We also analyzed the overlap in activity time of the three bat species using a non-parametric approach based on a kernel density estimated with the function density in the R package *Stats* 4.3.2. We obtained estimates of the Weitzman Superposition Coefficient Δ4 and its 95% confidence intervals using the *R* package *Overlapp* 0.3.9 (Meredith et al. 2024). We used the Watson (*U*² statistic) and Watson-Wheeler tests (*W* statistic) available in R package *Circular* 0.5-0 (Agostinelli and Lund 2023) to evaluate the goodness-of-fit of the activity time data for individual species, and to compare the activity time data distributions across pairs of bat species. Finally, we plotted the kernel density-time curves with default smoothing parameters of the density function with the R package *ggplot2* (Wickham 2016).

## RESULTS

### Dietary richness and composition

A total of 65 MOTUs were identified in the feces of the three bat species. We assigned 93.4–94.8% of MOTUs to order level, 77.9–83.8% to family/sub-family level, and 17.7– 30.9% to species level (Supplementary Data SD3 and SD4). Diets included 50 MOTUs from 71 *Molossus molossus*, 31 MOTUs from 44 *Myotis nigricans*, and 22 MOTUs from 21 *Neoeptesicus furinalis*. Estimated prey richness were 84.5 for *M. molossus*, 37.4 for *M. nigricans*, and 30.6 for *N. furinalis*. Sampling completeness at the observed sample sizes was approximately 88% for *M. molossus*, 91% for *M. nigricans*, and 85% for *N. furinalis* (Supplementary Data SD5).

Diet composition of the three bat species differed significantly at the order level (Pseudo-*F*_2, 133_ = 4.83, *P*-perm = 0.001). Ten insect orders were identified in the fecal samples, with nine in *M. molossus*, seven in *M. nigricans*, and five in *N. furinalis*.

Significant differences in the composition of insect orders occurred between *M. molossus* vs. *M. nigricans* (Pseudo-*t* = 2.52, *P*-perm = 0.001) and *M. molossus* vs. *N. furinalis* (Pseudo-*t* = 2.31, *P*-perm = 0.003). For *M. molossus*, the most frequently detected prey were Coleoptera, followed by Lepidoptera and Diptera. In contrast, *M. nigricans* had a higher incidence of Diptera, followed by Coleoptera and Hemiptera; whereas *N. furinalis* showed the highest incidence of Coleoptera, followed by Diptera and Hemiptera (Supplementary Data SD3; Fig. 2A).

**Fig. 2.**
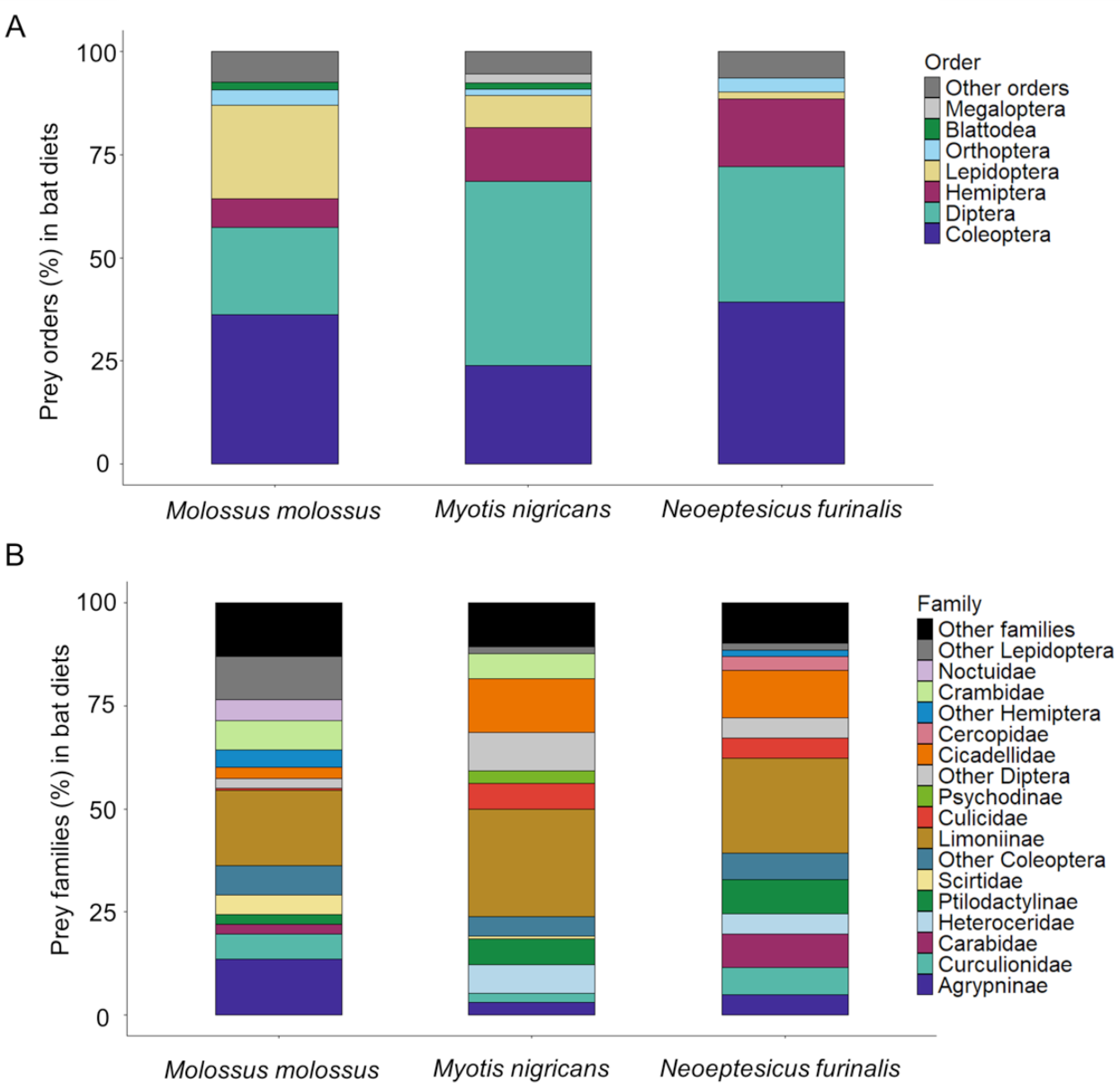
Percentage of prey in the diet of three species of aerial-hawking bats, classified by (A) order, and (B) family.

At the family/sub-family level, differences were also notable among bat species (Pseudo-*F*_2, 133_ = 3.92, *P*-perm = 0.001). Forty insect families/sub-families were identified in the fecal samples, with 32 in *M. molossus*, 19 in *M. nigricans*, and 13 in *N. furinalis*.

As above, the most notable differences in composition of insect families/sub-families were observed between *M. molossus* vs. *M. nigricans* (Pseudo-*t* = 2.39, *P*-perm = 0.001) and *M. molossus* vs. *N. furinalis* (Pseudo-*t* = 1.85, *P*-perm = 0.002). Limoniinae was the most frequently detected sub-family in the three bat species; however, *M. molossus* showed a higher incidence of Agrypninae, *N. furinalis* of Carabidae, and both *M. nigricans* and *N. furinalis* shared a greater incidence of Cicadellidae (Fig. 2B). All these insect taxa were collected in the rice fields (Supplementary Data SD3).

### Prey types

#### Sclerotization

The primary components in the diet of the two species with the strongest bites, *M. molossus* and *N. furinalis*, were prey with a high degree of sclerotization (Fig. 3A). Model selection supported bite force as the best predictor (wi = 0.70), with lower bite force in *M. nigricans* associated with a significantly lower incidence of strongly sclerotized prey in its diet (β = –0.52, *P*-value = 0.011). Nonetheless, the frequency of prey with intermediate sclerotization was similar across the three species (Fig. 3A). On the other hand, model selection showed similar support for species and bite force (wi = 0.51 and 0.49, respectively), with lower bite force in *M. nigricans* significantly increasing the presence of less-sclerotized prey (β = 0.62, *P*-value = 0.001), although this type of prey was also present in the diets of the other species with stronger bite force (Fig. 3A) (Supplementary Data SD6).

**Fig. 3.**
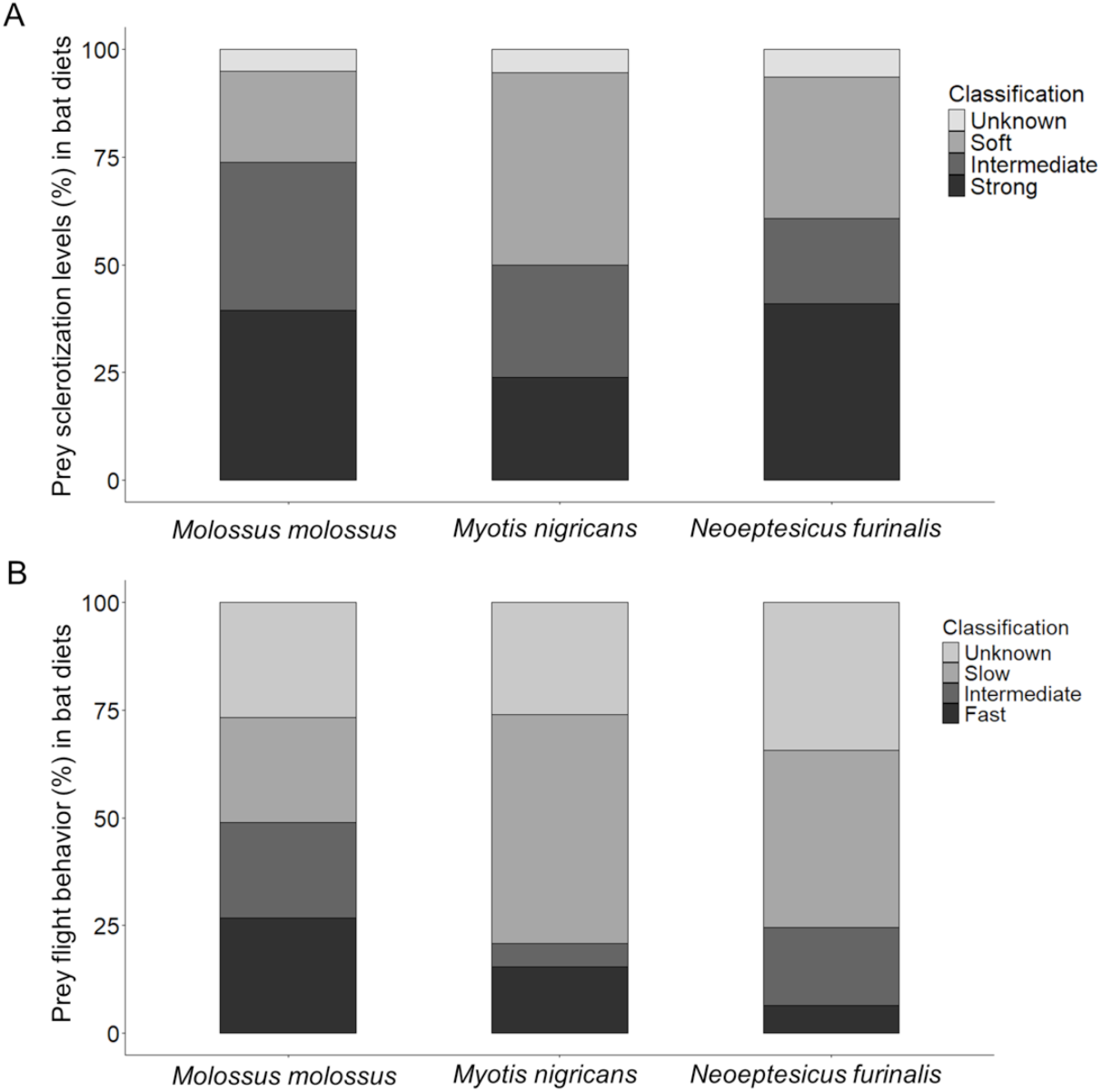
Percentage of prey in the diet of three species of aerial-hawking bats, classified by (A) degree of sclerotization, and (B) flight behavior.

#### Flight speed

Prey flight speed appeared to be closely associated with species that forage in uncluttered spaces, such as *M. molossus*. Fast-flying prey made up a major component of the diet of *M. molossus*, whereas in the other species their contribution was lower but not negligible (Fig. 3B). Model selection moderately supported functional group as the best predictor (wi = 0.38), with background-cluttered space foragers, *M. nigricans and N. furinalis*, associated with a significantly lower incidence of fast flyers prey in their diet (β = –0.78, *P*-value = 0.002). Model selection supported species as the best predictor of intermediate-flight prey (wi = 0.89), with *M. nigricans* showing a significantly lower presence of these prey in its diet (β = –1.19, *P*-value = 0.015). For this same prey type, functional group also had a negative effect (β = –0.87, *P*-value = 0.002), indicating that background-cluttered space foragers overall were less likely to capture intermediate flyers (Fig. 3B). On the other hand, model selection supported again functional group as the best predictor of slow-flying prey consumption (wi = 0.58), with background-cluttered space foragers showing a significantly higher incidence of slow flyers prey in their diets (β = 0.68, *P*-value = 0.00001). However, slow-flying prey were also present in the diet of *M. molossus* (Fig. 3B) (Supplementary Data SD6).

### Trophic niche breadth and overlap of IB species

Trophic niche breadth was highest in *M. molossus* (q₁=23.3), followed by *M. nigricans* (q₁=15.8) and *N. furinalis* (q₁=14.7). Pairwise niche overlap was highest between *N. furinalis* and *M. nigricans* (q₁=0.73), and lower between *M. molossus* and either *N. furinalis* (q₁=0.609) or *M. nigricans* (q₁=0.592) (Fig. 4; Supplementary Data SD5).

**Fig. 4.**
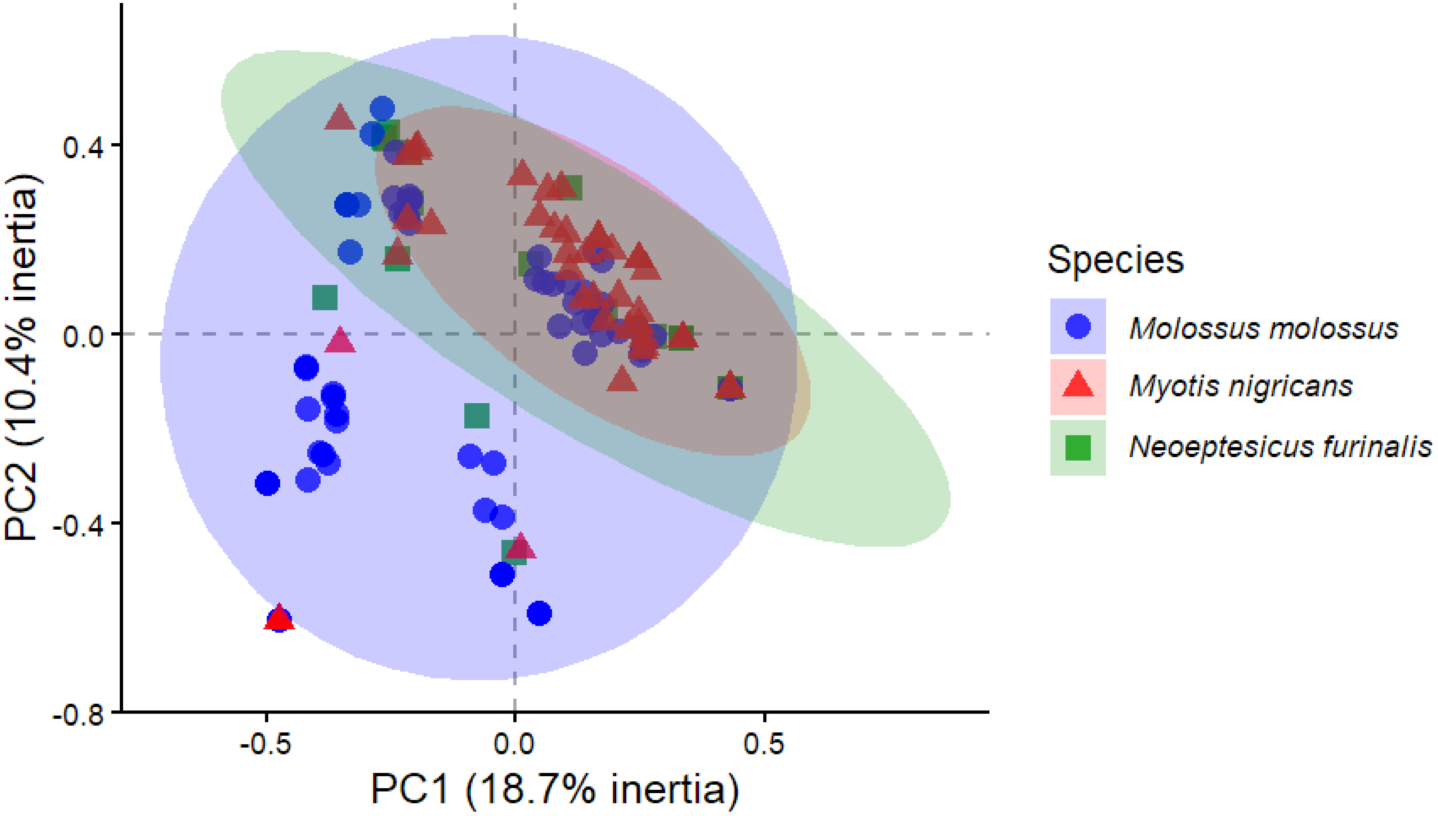
Hellinger-transformed PCA of dietary incidence (presence–absence of 65 MOTUs) in three species of aerial-hawking bats. Pterygota was excluded from the analyses, reducing the MOTU count by one (64 MOTUs).

### Bat activity time

*Molossus molossus* and *M. nigricans* represented 55.20% and 41.97%, respectively, of the sound files obtained in both farms (n = 6007). In contrast, *N. furinalis* had the lowest number of sound files during the entire study period (2.83%). The distribution of activity over time is not uniform for any of the species (*U*² = 11.1 to 241.8, all *P* < 0.01). The activity time of *M. molossus* did not overlap much with either *M. nigricans* (Δ4=0.36) or *N. furinalis* (Δ4=0.39). A high overlap in activity time between *N. furinalis* and *M. nigricans* was observed throughout the study (Δ4=0.93). However, peak activity times differed among the three species: 6:45 p.m. for *M. molossus*; 7:19 p.m. and 8:35 pm for *N. furinalis*; and 7:30 pm and 8:24 p.m. for *M. nigricans* (Fig. 5A-C). Additional tests comparing the distributions of activity over time in terms of orientation showed differences between *M. molossus* and both *M. nigricans* and *N. furinalis* (*W* = 248.1 to 2312.4, all *P* = 2.2e-16). In contrast, the same test comparing *M. nigricans* vs. *N. furinalis* was not significant (*W* = 3.1, *P* = 0.2).

**Fig. 5.**
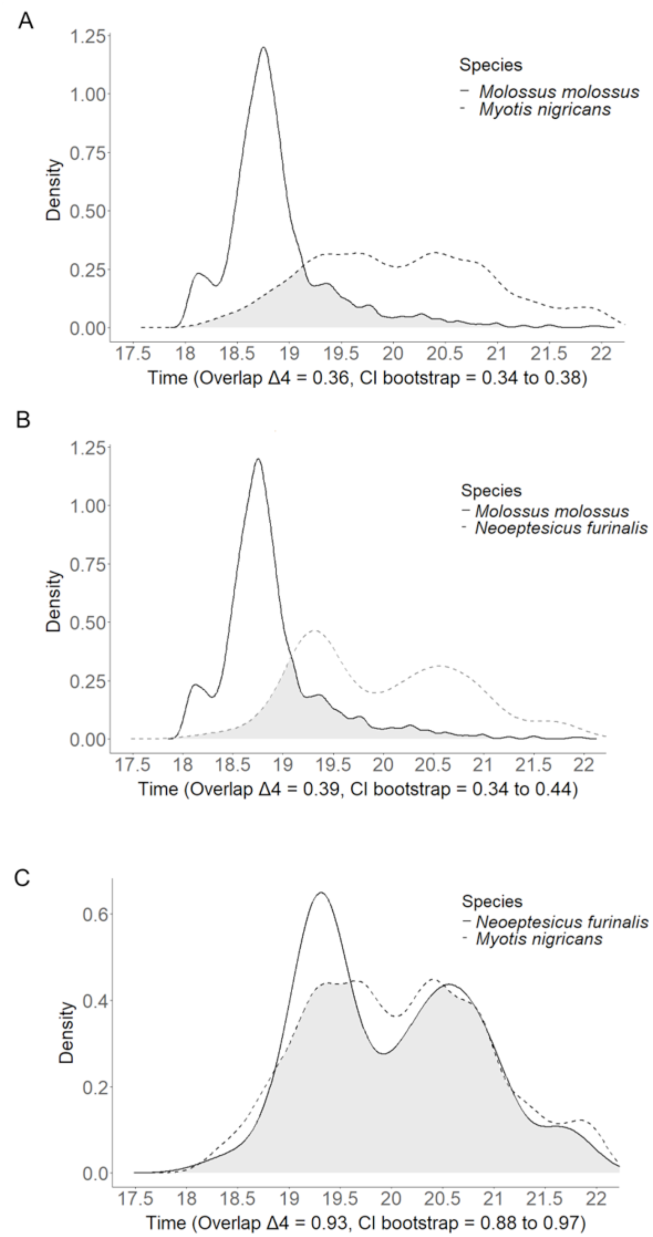
Time-density curves of bat species pairs with gray-shaded overlap areas, based on data from both farms. Bootstrap estimates and 95% confidence intervals for the overlap coefficient, Δ, are presented on the axis-x. The overlap coefficient Δ4 is used for sizes greater than 75.

## DISCUSSION

We investigated the diet and activity times of three species of aerial-hawking bats— *Molossus molossus*, *Neoeptesicus furinalis*, and *Myotis nigricans*—in rice fields using DNA metabarcoding of fecal samples and bioacoustics methods. These species exhibited distinct degrees of differentiation in dietary taxonomic richness and composition, biological attributes of prey consumed, and activity times. Consequently, they differed in the breadth and degree of overlap of their trophic niches, all of which resulted from traits related to bite force, hunting abilities, and intrinsic characteristics of each IB species. In addition to the traits analyzed, the proximity of roosts to foraging sites also influenced prey type by facilitating access to the insect community present in the rice fields. The humid conditions and high prey richness in rice fields therefore attractants of ecomorphologically distinct species, capable of exploiting these anthropogenic wetlands in distinct ways. These adaptive differences promote the coexistence of the three species, which appear to be structured by fine-scale trophic niche partitioning.

### Taxonomic richness and composition of prey

The diets of the IB species studied reflect the typical taxonomic richness and composition of insect communities inhabiting rice fields (Acosta et al. 2017; Yunus et al 2022). These rice fields undergo cyclical disturbances—such as flooding and harvesting—that promote temporal insect turnover, benefiting IB species that forage in such dynamic habitats. This may explain the high prey richness observed in the diets of the studied IB species—*M. molossus*, for example, showed nearly three times more prey families than other molossid bats from less diverse systems such as sugarcane or agro-urban environments (Bohmann et al. 2011; Aguiar et al. 2021). However, Kolkert et al. (2020) reported much higher prey richness—7.4 to 10.4 times more—in vespertilionid bat diets from subtropical agricultural regions with cotton crops in New South Wales.

This contrast likely reflects the broader range of habitat types present in that study, including forest, floodplain woodland and semi-arid grassland. By comparison, the rice fields in our study are embedded in a highly fragmented landscape, yet they appear to provide sufficient prey to buffer IB bats from the typical effects of habitat fragmentation.

Several insect families/sub-families detected in the IB diets, such as Limoniinae, Agrypninae, Curculionidae, Carabidae, Heteroceridae, Cicadellidae, and Crambidae, exhibit a broad range of life histories, developing in soil, water, or vegetation. Many of these insects engage in aerial activity during reproduction (Jackson and Campbell 1975; Pathak and Khan 1994; Traugott et al. 2015), which bats specialized in open and edge spaces are able to exploit. As expected, *M. molossus* showed the highest dietary taxonomic richness, consistent with its superior hunting abilities in these environments (Azofeifa et al. 2019). Moreover, edge-space species such as *M. nigricans* and *N. furinalis* appear to benefit from foraging in rice fields rather than depending on the few remaining forest patches, even if it means compromising prey richness in their diets. This opportunistic behavior underscores the importance of rice fields as key foraging habitats for IB in fragmented landscapes (Aizpurua et al. 2018).

### Prey types, trophic breadth, and overlap

Prey type was associated with hunting abilities, bite force, and intrinsic characteristics of each IB species. As predicted, *M. molossus*—adapted for fast, efficient flight in open habitats like the studied rice fields (Voigt and Holderied, 2012)—had a diet primarily composed of fast- and intermediate-flying insects (e.g., moths, beetles), while the two vespertilionids showed a higher representation of slow-flying insects such as leafhoppers and some dipterans. Tooth morphology, which ranges from compact in *M. nigricans* and *N. furinalis* to more robust in *M. molossus* (Mies et al. 1996; Loureiro et al. 2018; Novaes et al. 2022), may reflect a gradient in the contribution of soft- to hard-bodied prey (Freeman 1998), although, we did not detect this differentiation between *M. molossus* and the vespertilionid species. Instead, *M. nigricans* diverged more clearly from *N. furinalis* in the degree of prey sclerotization, likely due to its bite force being almost six times lower than in the latter species (Aguirre et al. 2002). Bite force therefore offered a more plausible explanation for the differences observed. However, although *M. molossus* has a slightly larger skull and somewhat higher bite force than *N. furinalis*, both species share a reduced third upper molar (Aguirre et al. 2002; Martínez-Coronel and Hortelano-Moncada 2020; Mies et al. 1996), which may enhance grinding and account for the similar representation of hard-bodied prey in their diets (Freeman 1998). Therefore, even though skull structure and bite force probably represent the most important drivers of hard-bodied prey processing in these bats (Giacomini et al. 2022), some dental could still play a role on it.

Despite the rich insect community present in these artificial wetlands, we were able to demonstrate that trophic niches are constrained by species-specific ecomorphological limitations (Mayberry et al. 2020). For example, although soft-bodied prey—mainly dipterans— are well represented in the diets of all three species, *M. nigricans* appears to specialize in them, which is consistent with its slow flight and relatively delicate skull. While this species can forage in open spaces by adjusting call bandwidth and duration to improve prey detection (Siemers et al. 2001; Bader et al. 2015), its biomechanical constraints likely limit the range of prey it can efficiently exploit. These results also support our prediction that *M. molossus* would include a broader range of prey in its diet, highlighting its trophic flexibility. This is further evidenced by the 24% of individuals (17/71) consuming prey detected once, compared to 11% in M. nigricans (5/44) and 10% in *N. furinalis* (2/21). Although singleton detections should be interpreted with caution due to the limitations of DNA metabarcoding (Alberdi and Gilbert 2019), these observations align with the idea that some individuals in generalist species may differ markedly in prey use (Bolnick et al. 2003), a trend that is receiving increasing support in bats (Arrizabalaga-Escudero et al. 2019; Magalhães de Oliveira et al. 2020).

In general, our analyses support *M. molossus* as the dominant forager, with low to moderate overlap in diet and activity time, which begins and peaks earlier than in the other two species. This molossid began hunting shortly before sunset (but see Esbérard and Bergallo 2010), coinciding with the rise in insect activity and the presence of some diurnal raptors—an ecological compromise that slow-flying bats would not efficiently cope with. The two vespertilionid species coexist with high overlap in both diet and activity times, although their activity peaks differ— similar to observations of other IB species reported in previous studies (Lambert et al. 2018; Beilke et al. 2021). These observations align with Aguiar and Antonini’s (2008) findings of high dietary overlap between these species based on morphological fecal analysis from the Brazilian Cerrado, and reinforce the idea that subtle ecological differences can facilitate coexistence even when resource use broadly overlaps. We believe that coexistence is possible despite higher trophic niche overlap, probably as a consequence of the high productivity of rice fields in artificial wetlands, where elevated prey abundance represents an almost unlimited food resource, leading to low competition and high trophic niche overlap (Krüger et al. 2012).

Comparisons of niche overlap across studies should be made with caution due to limitations in interpreting dietary data, particularly when using DNA metabarcoding. As Alberdi and Gilbert (2019) pointed out, Hill numbers offer a more robust framework for assessing diet diversity and overlap in metabarcoding studies. Direct comparisons using Pianka’s index are therefore not appropriate, as they may overestimate overlap.

Nevertheless, growing molecular evidence continues to reveal trophic flexibility in bats. Reported values of Pianka’s index have ranged from 0.50 to 0.83, in both cryptic species (Razgour et al. 2011) and inter- or intra-guild comparisons (Salinas-Ramos et al. 2015; Gordon et al. 2019; Segura-Trujillo et al. 2022).

### Concluding remarks

The coexistence between *M. molossus*, *M. nigricans*, and *N. furinalis* in the rice fields studied in the Venezuelan Llanos possibly reflects a combination of high insect availability in these artificial wetlands and partial differences in diet and activity time. *Molossus molossus* shows a higher incidence of fast-flying, medium-to-highly sclerotized prey, and its activity times started and peaked significantly earlier than in the other two bat species. Such divergence across these two trophic niche dimensions should reduce inter-specific competition and enhance coexistence between *M. molossus* and the vespertilionids, which agree with niche theory predictions (Finke and Snyder 2008). *Myotis nigricans* and *N. furinalis* are less specialized for foraging in open environments, and their trophic niches are constrained by species-specific ecomorphological limitations. Their diets were primarily composed of slow-flying prey and showed high temporal overlap in activity, though with distinct peak times. A stronger bite in *N. furinalis*, however, may have allowed for the inclusion of more strongly sclerotized prey than in *M. nigricans*, suggesting that differences in prey hardness reduce competition despite similar flight strategies. Although our analyses did not account for seasonal or cyclical variation in rice cultivation, the non-migratory nature and year-round presence of these species in the study area suggest that trophic interactions may remain relatively stable over time. In this context, fine-scale trophic niche partitioning—facilitated by the humid, prey-rich environment of rice fields—appears to mediate the coexistence of IB species with marked ecomorphologically differences.

## Acknowledgements

Special thanks to José Antonio González-Carcacía (IVIC) for his support during some fieldwork sessions. Claudia Domínguez (IVIC) and Abimel Moreno (IVIC) provided assistance in identifying collected insects. Delphine Rioux (LECA) and Eric Coissac (LECA) assisted with molecular and bioinformatics analyses.

## Supplementary Data

Supplementary Data SD1. — Taxonomic identification notes and ecomorphological traits of the IB species

Supplementary Data SD2. — Prey hardness (order level) and flight type (family level) of dietary items in three IB species.

Supplementary Data SD3. — Percentage of prey by taxon (Pi) in the diet of three species of aerial hawking bats at North-western Venezuela’s rice crops.

Supplementary Data SD4. —Sequences of insect MOTUs (Inse01_motus and Arth02_motus) used for the assignment of MOTUs in bat feces (bat_feces_Inse01.motus and bat_feces_Arth02_motus).

Supplementary Data SD5. — Estimated diversity, trophic niche breadth and dietary overlap for three IB species.

Supplementary Data SD6— GLMs assessing the impact of bat factors (species, functional group, and bite force) on prey characteristics.

