## Supplementary material for "Trophic flexibility favors coexistence of three aerial-hawking bat species in Venezuelan rice fields": Supp.File 1

Yara Azofeifa Romero (1, 2, †)
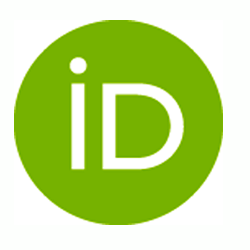
**<https://orcid.org/0000-0003-2874-4207>**, Jafet M Nassar (1)
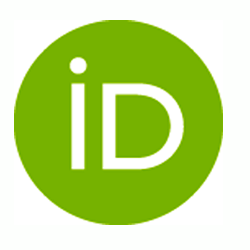
**<https://orcid.org/0000-0001-9808-9362>**, Jesús Mavárez (3, ‡)
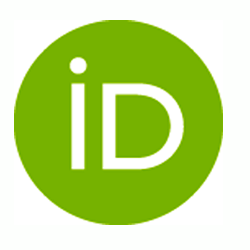
**<https://orcid.org/0000-0002-2483-0178>**

**Notes on the taxonomic identification of the three IB species**

During the 2013–2014 fieldwork, we used the keys of Linares (1998) and Gardner (2007). Later, while preparing this manuscript, we verified taxonomic changes using the key of Días et al. (2016), the Mammal Diversity Database (Burgin et al. 2025), and scientific publications such as González-Ruiz et al. (2011), Novaes et al. (2017), and Ramírez-Chaves et al. (2021).

Species similar to *Molossus molossus* include *M. coibensis and M. alvarezi;* however, *M. coibensis* is smaller and *M. alvarezi* is larger than *M. molossus*. The fur of *M. molossus* was lighter in color than that of the other two species. Overall, this species is very common and easy to identify.

*Neoeptesicus furinalis* could be confused with *N. orinoensis* and *N. brasiliensis*; however, *N. orinoensis* is smaller and *N. brasiliensis* is larger than *N. furinalis*. A distinguishing dental feature of *N. furinalis* is its well-developed upper first incisor, which bears a secondary cusp. Additionally, the upper premolar is located adjacent to the canine and reaches more than half its height.

*Myotis nigricans* and *M. riparius* are similar in size; however, the fur of *M. nigricans* is silky with a more uniform coloration, whereas the latter species has woolly fur with a distinct bicolored ventral pattern.

**Ecomorphological traits for the three bat species from literature**

*Neoeptesicus furinalis* has a body weight that is 54% (or 0.54 times) of *M. molossus* and 168% (or 1.68 times) of that of *Myotis nigricans*, indicating an intermediate mass. Its forearm length is 99% that of *M. molossus* and 112% (or 1.12 times) that of *M. nigricans*, indicating similar forearm proportions to *M. molossus*. Cranial length in *N. furinalis* is 93% of *M. molossus* and 116% (or 1.16 times) that of *M. nigricans*, indicating an intermediate skull length. The bite force of *N. furinalis* is 87.5% that of *M. molossus* and approximately 575% (or 5.75 times) that of *M. nigricans*, indicating a bite force comparable to the M. molossus. In terms of flight-related parameters, *N. furinalis* and *M. nigricans,* both aerial insectivorous that forage in *background-cluttered space,* show lower wing loading and aspect ratio than *M. molossus*, an aerial insectivorous adapted to uncluttered space (Table SD1a).

**Table SD1a.** Reference values of ecomorphological traits related to trophic niche differentiation among three IB species. Data sources: Linares (1998) for weight and forearm length; Wilson and LaVal (1974), Mies et al. (1996), Loureiro et al. (2018) for skull length; Mies et al. (1996), Loureiro et al. (2018), Novaes et al. (2022) for length of maxillary toothrow and number of teeth; Aguirre et al. (2002) for bite force; Amador et al. (2020) for wing loading and aspect ratio.

|  | Molmol | Neofur | Myonig |
| --- | --- | --- | --- |
| Average weight (g) | 14.4* | 7.8* | 4.65* |
| Average forearm length (mm) | 38.5* | 38.2* | 34.1* |
| Average skull length (mm) | 17.3 | 16.1 | 13.9 |
| Length of maxillary toothrow (mm) | 6.3 | 5.4 | 5.0 |
| Number of teeth | 26.0 | 32.0 | 38.0 |
| Max. bite force (N) | 8.34 | 7.3 | 1.27 |
| Wing loading (Pa) | 14.1 | 7.3 | 6.1 |
| Aspect ratio | 9.2 | 6.2 | 6.5 |

*Values marked with an asterisk (*) represent averages estimated from the minimum and maximum values reported by Linares (1998) in Venezuela.*

**Forearm and body mass measurements of individuals analyzed for diet**

In this study, *N. furinalis* has a body weight that is 56.8% (or 0.57 times) that of *M. molossus* and 166.5% (or 1.67 times) that of *M. nigricans*, indicating an intermediate mass. Its forearm length is 94.6% that of *M. molossus* and 110.5% (or 1.11 times) that of *M. nigricans*, suggesting similar forearm proportions to M. molossus (Table SD1b).

**Table SD1b.** Forearm and body mass measurements of individuals analyzed for diet.

| **LECA code** | **Species** | **Body weight (g)** | **Forearm length (mm)** |
| --- | --- | --- | --- |
| F006 | Molmol | 12.0 | 40.2 |
| F007 | Molmol | 13.1 | 38.75 |
| F008 | Molmol | 11.8 | 39.55 |
| F010 | Molmol | 15.2 | 39.85 |
| F012 | Molmol | 13.9 | 39.3 |
| F014 | Molmol | 13.6 | 41.1 |
| F015 | Molmol | 11.4 | 38.6 |
| F017 | Molmol | 11.1 | 41.7 |
| F018 | Molmol | 13.9 | 40.15 |
| F019 | Molmol | 12.1 | 39.4 |
| F020 | Molmol | 11.9 | 41.55 |
| F021 | Molmol | 12.2 | 39.5 |
| F022 | Molmol | 12.1 | 38.8 |
| F023 | Myonig | 3.8 | 31.7 |
| F024 | Myonig | 3.9 | 33.7 |
| F025 | Molmol | 11.4 | 38.9 |
| F042 | Molmol | 12.7 | 39.5 |
| F043 | Molmol | 11.1 | 40.95 |
| F044 | Molmol | 14.5 | 40.95 |
| F045 | Molmol | 12.2 | 39.2 |
| F046 | Molmol | 11.1 | 39.45 |
| F047 | Molmol | 12.4 | 40.3 |
| F048 | Neofur | 4.2 | 35.75 |
| F049 | Myonig | 4.3 | 34.3 |
| F050 | Neofur | 4.5 | 37.0 |
| F051 | Myonig | 3.6 | 34.25 |
| F054 | Myonig | 4.4 | 33.8 |
| F055 | Myonig | 3.8 | 33.2 |
| F056 | Molmol | 11.8 | 38.85 |
| F057 | Molmol | 12.6 | 39.45 |
| F058 | Molmol | 11.1 | 39.25 |
| F059 | Molmol | 12.3 | 39.35 |
| F060 | Molmol | 10.2 | 38.1 |
| F061 | Molmol | 15.7 | 41.5 |
| F062 | Molmol | 12.5 | 40.45 |
| F063 | Myonig | 5.1 | 32.5 |
| F064 | Myonig | 4.0 | 33.95 |
| F067 | Myonig | 4.3 | 32.75 |
| F068 | Molmol | 11.9 | 39.3 |
| F069 | Molmol | 11.9 | 39.75 |
| F070 | Molmol | 11.2 | 39.9 |
| F071 | Molmol | 14.4 | 39.7 |
| F072 | Molmol | 11.3 | 40.05 |
| F073 | Molmol | 12.0 | 38.75 |
| F094 | Molmol | 13.7 | 37.85 |
| F095 | Molmol | 14.3 | 40.15 |
| F097 | Molmol | 15.1 | 40.2 |
| F100 | Molmol | 12.3 | 37.7 |
| F102 | Myonig | 5.0 | 34.2 |
| F103 | Myonig | 3.4 | 32.2 |
| F105 | Neofur | 7.5 | 37.25 |
| F106 | Neofur | 10.1 | 38.25 |
| F107 | Myonig | 4.0 | 32.0 |
| F108 | Molmol | 12.6 | 40.75 |
| F109 | Molmol | 13.2 | 41.15 |
| F110 | Molmol | 12.3 | 39.0 |
| F111 | Molmol | 13.2 | 38.9 |
| F112 | Molmol | 12.3 | 39.8 |
| F114 | Molmol | 13.4 | 38.1 |
| F115 | Molmol | 12.5 | 40.25 |
| F118 | Myonig | 4.0 | 35.7 |
| F119 | Myonig | 5.7 | 35.0 |
| F120 | Myonig | 3.5 | 32.85 |
| F121 | Myonig | 4.9 | 34.2 |
| F122 | Myonig | 4.5 | 34.1 |
| F123 | Neofur | 6.1 | 36.95 |
| F124 | Neofur | 7.8 | 38.95 |
| F125 | Neofur | 7.2 | 37.5 |
| F126 | Neofur | 6.5 | 36.8 |
| F127 | Molmol | 12.2 | 39.9 |
| F128 | Molmol | 12.6 | 39.5 |
| F129 | Molmol | 13.8 | 41.05 |
| F130 | Molmol | 13.9 | 39.7 |
| F131 | Molmol | 13.3 | 40.2 |
| F133 | Myonig | 5.0 | 34.4 |
| F134 | Myonig | 4.2 | 34.7 |
| F136 | Myonig | 5.2 | 35.25 |
| F137 | Myonig | 5.2 | 36.0 |
| F138 | Myonig | 6.1 | 34.5 |
| F139 | Myonig | 4.4 | 34.3 |
| F140 | Myonig | 3.9 | 32.9 |
| F142 | Neofur | 9.9 | 37.1 |
| F143 | Neofur | 7.7 | 36.3 |
| F144 | Neofur | 7.2 | 37.2 |
| F145 | Neofur | 7.6 | 37.0 |
| F146 | Neofur | 7.9 | 39.5 |
| F147 | Molmol | 13.2 | 39.3 |
| F148 | Molmol | 11.7 | 39.2 |
| F149 | Molmol | 11.6 | 39.2 |
| F150 | Molmol | 14.0 | 39.1 |
| F151 | Molmol | 14.6 | 40.1 |
| F152 | Molmol | 12.2 | 40.4 |
| F153 | Molmol | 12.9 | 39.1 |
| F154 | Molmol | 14.4 | 39.2 |
| F155 | Myonig | 4.8 | 34.1 |
| F156 | Myonig | 4.5 | 34.0 |
| F157 | Myonig | 4.1 | 33.6 |
| F158 | Myonig | 4.2 | 34.2 |
| F159 | Myonig | 4.2 | 35.6 |
| F160 | Myonig | 3.9 | 32.5 |
| F162 | Molmol | 14.7 | 39.75 |
| F164 | Molmol | 14.0 | 39.5 |
| F165 | Molmol | 13.8 | 39.5 |
| F166 | Molmol | 15.4 | 39.3 |
| F167 | Molmol | 12.8 | 39.1 |
| F169 | Molmol | 11.8 | 39.7 |
| F171 | Neofur | 7.8 | 38.7 |
| F172 | Neofur | 7.5 | 36.8 |
| F173 | Neofur | 8.0 | 37.6 |
| F174 | Neofur | 6.5 | 35.9 |
| F175 | Neofur | 8.2 | 40.3 |
| F176 | Neofur | 7.9 | 37.2 |
| F177 | Myonig | 4.6 | 35.3 |
| F178 | Myonig | 3.3 | 31.9 |
| F179 | Myonig | 4.2 | 33.4 |
| F180 | Myonig | 4.8 | 33.5 |
| F181 | Neofur | 6.9 | 37.5 |
| F182 | Neofur | 7.2 | 36.8 |
| F183 | Myonig | 4.7 | 35.6 |
| F184 | Myonig | 4.3 | 34.5 |
| F185 | Myonig | 3.6 | 32.5 |
| F186 | Myonig | 5.2 | 33.4 |
| F187 | Myonig | 6.1 | 34.1 |
| F190 | Molmol | 13.1 | 38.75 |
| F192 | Molmol | 12.1 | 38.9 |
| F193 | Molmol | 12.8 | 38.6 |
| F194 | Myonig | 4.6 | 34.8 |
| F197 | Myonig | 4.0 | 35.3 |
| F198 | Myonig | 3.8 | 33.1 |
| F199 | Myonig | 4.7 | 33.5 |
| F200 | Myonig | 4.2 | 33.4 |
| F204 | Molmol | 15.1 | 40.0 |
| F205 | Molmol | 15.3 | 38.85 |
| F206 | Molmol | 14.1 | 39.0 |
| F207 | Molmol | 13.0 | 38.0 |
| F208 | Molmol | 16.6 | 39.5 |
