## Supplementary material for "Trophic flexibility favors coexistence of three aerial-hawking bat species in Venezuelan rice fields": Supp.File 2

Yara Azofeifa Romero (1, 2, †)
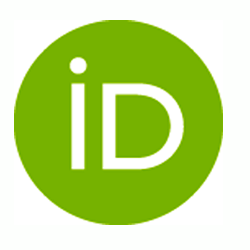
**<https://orcid.org/0000-0003-2874-4207>**, Jafet M Nassar (1)
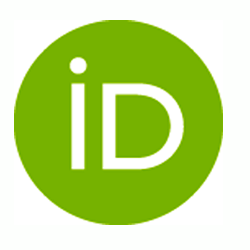
**<https://orcid.org/0000-0001-9808-9362>**, Jesús Mavárez (3, ‡)
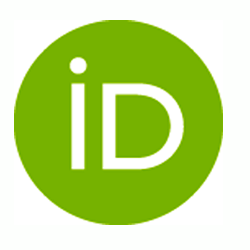
**<https://orcid.org/0000-0002-2483-0178>**

**Table SD2a.** Prey hardness at the order level for items in the diet of three IB species.

| **Taxa** | **Classification** |
| --- | --- |
| Coleoptera, and Hemiptera | Strongly sclerotized |
| Lepidoptera, Odonata, Orthoptera,  and Hemiptera | Medium sclerotized |
| Diptera | Less sclerotized |

**Tables SD2b.** Prey flight type at the family level for items in the diet of three IB species.

| **Taxa** | **Classification** | **Description** | **Reference** |
| --- | --- | --- | --- |
| Limoniidae | Slow flyers | Slow and awkward flights of long vertical range (up to 100 m) | Jackson DM, Campbell RL. 1975. Biology of the European crane fly, *Tipula paludosa* Meigen, in western Washington (Tipulidae; Diptera). Pullman (WA): Washington State University, College of Agriculture Research Center. Technical Bulletin 81. |
| Culicidae | Slow flyers | Slow flight with elliptic loops within swarms, although these can maneuver in the air with great agility | Gibson G. 1985. Swarming behaviour of the mosquito Culex pipiens quinquefasciatus: a quantitative analysis. *Physiological Entomology* 10(3):283–296. <https://doi.org/10.1111/j.1365-3032.1985.tb00049.x>  Lams SM. 2012. Free flight of the mosquito *Aedes aegypti*. A*rXiv,1205.5260.* <https://doi.org/10.48550/arXiv.1205.5260> |
| Drosophylidae | Fast flyers | Speed and optomotor control | David C. 1979. Optomotor control of speed and height by free-flying *Drosophila*. *The Journal of experimental biology* 82(1):389–392. <https://doi.org/10.1242/jeb.82.1.389> |
| Elateridae (Agrypninae) | Intermediate flyers | Infrequent flyers, they make short-range local movements. They perform vertical flights near or far up to 2 m. | Parker WE, Howard JJ. 2001. The biology and management of wireworms (*Agriotes* spp.) on potato with particular reference to the U.K.: Biology and management of wireworms on potato. *Agricultural and Forest Entomology* 3(2):85–98. <https://doi.org/10.1046/j.1461-9563.2001.00094.x> |
| Crambidae | Fast flyers | Efficient and maneuverable flyers, capable of reaching height > 50 m | Arrizabalaga-Escudero A, Merckx T, García-Baquero G, Wahlberg N, Aizpurua O, Garin I, Goiti U, Aihartza J. 2019. Trait-based functional dietary analysis provides a better insight into the foraging ecology of bats. *Journal of Animal Ecology* 88(10):1587–1600. <https://doi.org/10.1111/1365-2656.13055> |
| Noctuidae | Fast flyers | Strong flyers, capable of reaching heights > 50 m | Arrizabalaga-Escudero A, Merckx T, García-Baquero G, Wahlberg N, Aizpurua O, Garin I, Goiti U, Aihartza J. 2019. Trait-based functional dietary analysis provides a better insight into the foraging ecology of bats. *Journal of Animal Ecology* 88(10):1587–1600. <https://doi.org/10.1111/1365-2656.13055>  Lingren PD, Raulston JR, Popham TW, Wolf WW, Lingren PS, Esquivel JF. 1995. Flight behavior of corn earworm (Lepidoptera: Noctuidae) moths under low wind speed conditions. *Environmental entomology* 24(4):851–860. <https://doi.org/10.1093/ee/24.4.851> |
| Curculionidae | Intermediate flyers | At first, they have an uncontrolled flight, they take off with their wings, they then fly downwind for considerable distances.  They perform vertical flights of up to 30-50 m | Solbreck C. 1980. Dispersal distances of migrating pine weevils, *Hylobius abietis* (Coleoptera: Curculionidae). *Entomologia Experimentalis et Applicata* 28(2):123–131. <https://doi.org/10.1111/j.1570-7458.1980.tb02997.x> |
| Cicadellidae | Slow flyers | Poor flyers, perform horizontal and vertical flights.  They jump to start the flight. The most common jumps are the horizontal ones over the plant canopy. | Rodriguez CM, Madden LV, Nault LR. 1992. Diel flight periodicity of *Graminella nigrifrons* (Homoptera: Cicadellidae). *Annals of the Entomological Society of America* 85(6):792–798. <https://doi.org/10.1093/aesa/85.6.792>  Chancellor TCB, Cook AG, Heong KL, Villareal S. 1997. The flight activity and infectivity of the major leafhopper vectors (Hemiptera: Cicadellidae) of rice tungro viruses in an irrigated rice area in the Philippines. *Bulletin of Entomological Research* 87(3):247–258. <https://doi.org/10.1017/S0007485300037196>  Lopes JRS, Nault LR, Phelan PL. 1995. Periodicity of diel activity of *Graminella nigrifrons* (Homoptera: Cicadellidae) and implications for leafhopper dispersal. *Annals of the Entomological Society of America* 88(2):227–233. <https://doi.org/10.1093/aesa/88.2.227>  Clemente CJ, Goetzke HH, Bullock JMR, Sutton GP, Burrows M, Federle W. 2017. Jumping without slipping: leafhoppers (Hemiptera: Cicadellidae) possess special tarsal structures for jumping from smooth surfaces. *Journal of The Royal Society Interface* 14(130):20170022. <http://dx.doi.org/10.1098/rsif.2017.0022> |
| Delphacidae | Slow flyers | Weak flier, highly dependent on air currents for acceleration and long-distance travel | Rosenberg LJ, Magor JI. 1983. Flight duration of the brown planthopper, *Nilaparvata lugens* (Homoptera: Delphacidae). *Ecological Entomology* 8(3):341-350. <https://doi.org/10.1111/j.1365-2311.1983.tb00514.x> |
| Carabidae | Intermediate flyers | At first, they have an uncontrolled flight because they jump,  then they disperse without control over the height of the flight | Boiteau G, Bousquet Y, Osborn W. 2000. Vertical and temporal distribution of Carabidae and Elateridae in flight above an agricultural landscape. *Environmental Entomology* 29(6):1157–1163. <https://doi.org/10.1603/0046-225X-29.6.1157> |
| Hydrophilidae | Slow flyers | Poor flight ability | Jackson DJ. 1956. The capacity for flight of certain water beetles and its bearing on their origin in the Western Scottish Isles. *Proceedings of the Linnean Society of London* 167(1):76–96. <https://doi.org/10.1111/j.1095-8312.1956.tb00771.x> |
| Orthoptera | Fast flyers | Long-winged morphs are strong flyers | Jiang CJ, Zhang BC, Chen WF, Zhang QW, Zhao ZW, An CJ, et al. 2012. Variations in the ultrastructure of the flight muscles of the polymorphic cricket, *Gryllus firmus* (Orthoptera: Gryllidae). *European Journal of Entomology* 109(4):579–586. <https://doi.org/10.14411/eje.2012.072> |
