## Supplementary material for "Trophic flexibility favors coexistence of three aerial-hawking bat species in Venezuelan rice fields": Supp.File 3

Yara Azofeifa Romero (1, 2, †)
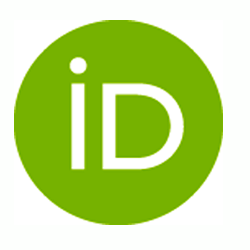
**<https://orcid.org/0000-0003-2874-4207>**, Jafet M Nassar (1)
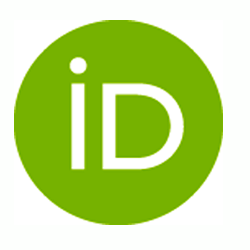
**<https://orcid.org/0000-0001-9808-9362>**, Jesús Mavárez (3, ‡)
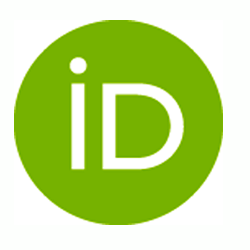
**<https://orcid.org/0000-0002-2483-0178>**

**Table SD3.** Percentage of prey by taxon (Pi) in the diet of *M. molossus*, *M. nigricans*, and *N. furinalis* at rice fields in Northwestern Venezuela. Pi = (∑Ni/N total) × 100, where ∑Ni is the total number of prey items for taxon i found in the diet and N total is the total number of prey items across all taxa in the diet. Fecal samples from the three species were collected between November 2013 and September 2014. * Collected in the studied rice fields (Moreno et al. 2020).

| Taxa | *Molossus molossus*  N _BATS_ = 71 N _PREYS =_ 213 | *Myotis nigricans*  N _BATS_ = 44 N _PREYS_ = 130 | *Neoeptesicus furinalis* N _BATS_ = 21  N _PREYS_ = 61 |
| --- | --- | --- | --- |
| **Order Coleoptera** |  |  |  |
| Suborder Adephaga |  |  |  |
| *Carabidae |  |  |  |
| Unknown Carabidae | 2.35 | 0.00 | 6.56 |
| Cicindelinae | 0.00 | 0.00 | 1.64 |
| *Dytiscidae | 0.47 | 0.00 | 0.00 |
| Suborder Polyphaga |  |  |  |
| Unknown Polyphaga | 2.35 | 1.54 | 3.28 |
| Unknown Cucujiformia | 0.94 | 0.00 | 0.00 |
| *Elateridae: Agrypninae (*Aeolus* sp) | 13.62 | 3.08 | 4.92 |
| *Ptilodactylinae | 2.35 | 6.15 | 8.20 |
| *Curculionidae |  |  |  |
| Unknown Curculionidae | 0.47 | 0.00 | 1.64 |
| *Scolytinae | 0.00 | 0.77 | 0.00 |
| *Erirhininae*: Lissorhoptrus sp* | 5.63 | 1.54 | 4.92 |
| *Scirtidae | 4.69 | 0.77 | 0.00 |
| *Scarabaeidae: Dynastinae | 0.47 | 0.00 | 0.00 |
| *Hydrophilidae | 0.94 | 1.54 | 0.00 |
| *Heteroceridae | 0.00 | 6.92 | 4.92 |
| *Tenebrionidae | 0.47 | 0.00 | 1.64 |
| *Chrysomelidae | 0.94 | 0.00 | 1.64 |
| Throscidae | 0.00 | 0.77 | 0.00 |
| *Zopheridae: Colydiinae | 0.47 | 0.77 | 0.00 |
| **Order Diptera** |  |  |  |
| Unknown Diptera | 0.47 | 0.00 | 1.64 |
| Suborder Nematocera |  |  |  |
| Unknown Nematocera | 0.00 | 1.54 | 0.00 |
| Sciaroidea | 0.00 | 1.54 | 1.64 |
| *Tipuloidea: Limoniinae | 18.31 | 26.15 | 22.95 |
| Tipuloidea: Pediciidae | 0.47 | 0.00 | 0.00 |
| *Culicoidea: Culicidae | 0.47 | 6.15 | 4.92 |
| Chironomoidea: Ceratopogonidae | 0.47 | 0.00 | 0.00 |
| Psychodoidea: Psychodinae | 0.00 | 3.08 | 0.00 |
| Suborder Brachycera |  |  |  |
| Unknown Muscomorpha | 0.00 | 1.54 | 0.00 |
| Calyptratae | 0.47 | 0.00 | 0.00 |
| Cyclorrhapha | 0.00 | 3.08 | 1.64 |
| Drosophilidae | 0.47 | 0.77 | 0.00 |
| Stratyiomidae | 0.00 | 0.77 | 0.00 |
| **Order Lepidoptera** |  |  |  |
| Unknown Lepidoptera | 0.47 | 0.00 | 0.00 |
| Suborder Glossata |  |  |  |
| Rhopalocera | 2.35 | 0.00 | 0.00 |
| Obtectomera | 6.57 | 0.77 | 0.00 |
| *Crambidae |  |  |  |
| Unknown Crambidae | 2.35 | 3.85 | 0.00 |
| **Rupela albinella* | 4.23 | 1.54 | 0.00 |
| *Diatraea saccharalis* | 0.47 | 0.00 | 0.00 |
| *Eoreuma loftini* | 0.00 | 0.77 | 0.00 |
| *Noctuidae |  |  |  |
| Unknown Noctuidae | 0.94 | 0.00 | 0.00 |
| **Spodoptera frugiperda* | 3.29 | 0.00 | 0.00 |
| *Leucania*sp. | 0.47 | 0.00 | 0.00 |
| *Condica sutor* | 0.47 | 0.00 | 0.00 |
| Hesperiidae: Eudamidae | 0.00 | 0.00 | 1.64 |
| Geometridae | 0.00 | 0.77 | 0.00 |
| Saturnidae | 0.47 | 0.00 | 0.00 |
| Erebidae | 0.47 | 0.00 | 0.00 |
| **Order Orthoptera** |  |  |  |
| Unknown Orthoptera | 1.88 | 0.77 | 3.28 |
| Suborder Ensifera |  |  |  |
| *Gryllidae | 1.88 | 0.77 | 0.00 |
| **Order Hemiptera** |  |  |  |
| Unknown Hemiptera | 0.94 | 0.00 | 0.00 |
| Suborder Heteroptera |  |  |  |
| Panheteroptera | 0.47 | 0.00 | 0.00 |
| *Belostomatidae | 0.94 | 0.00 | 1.64 |
| *Lygaeidae | 0.47 | 0.00 | 0.00 |
| *Pentatomidae: *Oebalus ypsilongriseus* | 0.47 | 0.00 | 0.00 |
| Suborder Auchenorrhyncha |  |  |  |
| *Cicadellidae |  |  |  |
| Unknown Cicadellidae | 1.41 | 2.31 | 3.28 |
| **Graminella* sp | 1.41 | 10.77 | 8.20 |
| *Cercopidae | 0.00 | 0.00 | 3.28 |
| *Delphacidae: *Tagosodes orizicolus* | 0.94 | 0.00 | 0.00 |
| **Order Himenoptera**: Formicidae | 1.41 | 0.00 | 0.00 |
| **Order Blattodea** |  |  |  |
| Blaberidae | 0.47 | 0.00 | 0.00 |
| Rhinotermitidae | 0.47 | 1.54 | 0.00 |
| Kalotermitidae | 0.94 | 0.00 | 0.00 |
| **Order Neuroptera**: *Chrysopinae | 0.47 | 0.00 | 0.00 |
| **Order Megaloptera**: Corydalidae | 0.00 | 2.31 | 0.00 |
| **Order Odonata**: Coenagrionidae | 0.47 | 0.00 | 0.00 |
| **Unknown Pterygota** | 5.16 | 5.38 | 6.56 |
| Total (%) | 100 | 100 | 100 |
| Mean | 1.54 | 1.54 | 1.54 |
| Median | 0.47 | 0.00 | 0.00 |
| 2sd | 2.99 | 3.70 | 3.44 |
| Min | 0.00 | 0.00 | 0.00 |
| Max | 18.31 | 26.15 | 22.95 |
| 25th percentile | 0.47 | 0.00 | 0.00 |
| 75th percentile | 1.41 | 1.54 | 1.64 |
