## Supplementary material for "Trophic flexibility favors coexistence of three aerial-hawking bat species in Venezuelan rice fields": Supp.File 5

Yara Azofeifa Romero (1, 2, †)
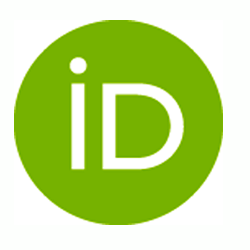
**<https://orcid.org/0000-0003-2874-4207>**, Jafet M Nassar (1)
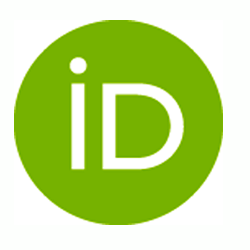
**<https://orcid.org/0000-0001-9808-9362>**, Jesús Mavárez (3, ‡)
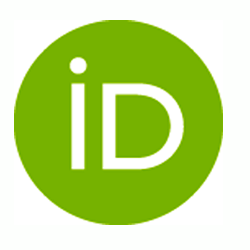
**<https://orcid.org/0000-0002-2483-0178>**

**Table SD5a**. Estimated diversity values (Hill numbers q = 0, 1, 2) for three IB species based on DNA metabarcoding data**.** Observed values and extrapolated estimators are shown for species richness (q = 0), Shannon diversity (q = 1), and Simpson diversity (q = 2), with associated standard errors (s.e.) and 95% confidence intervals (LCL–UCL).

| IB Species | q | Observed | Estimator | s.e. | LCL | UCL |
| --- | --- | --- | --- | --- | --- | --- |
| Molmol | 0 | 49 | 84.49296 | 19.35947 | 49 | 122.4368 |
| Molmol | 1 | 23.33 | 29.00994 | 2.725675 | 23.66771 | 34.35216 |
| Molmol | 2 | 12.863808 | 13.54522 | 1.363594 | 10.87262 | 16.21781 |
| Neofur | 0 | 21 | 30.64286 | 12.54248 | 21 | 55.22567 |
| Neofur | 1 | 14.682404 | 19.3823 | 3.020147 | 13.46292 | 25.30168 |
| Neofur | 2 | 10.058824 | 11.63266 | 1.999429 | 7.713847 | 15.55146 |
| Myonig | 0 | 30 | 37.39063 | 12.91909 | 30 | 62.71157 |
| Myonig | 1 | 15.821024 | 18.60266 | 1.681335 | 15.3073 | 21.89801 |
| Myonig | 2 | 8.925664 | 9.405318 | 0.988691 | 7.467519 | 11.34312 |

* Pterygota were excluded from the analyses, reducing the MOTU count by one. As a result, in the following analyses, the diets comprised 49 MOTUs from 71 *M. molossus* (Molmol) samples, 30 MOTUs from 44 *M. nigricans* (Myonig) samples, and 21 MOTUs from 21 *N. furinalis* (Neofur) samples.

**Table SD5b**. Summary of sampling information and prey detection frequencies per bat species. T= Number of fecal simples; U= Total number of times (incidences) a prey item was detected in the samples; S.obs= Observed MOTU richness; SC= Sampling Coverage (Sampling completeness); Q1-Q10= Number of MOTUs detected in exactly 1 to 10 samples, respectively. Only Q1 (singletons) is used in the estimation of sampling coverage (SC), as a measure of the proportion of unobserved diversity.

| Specie | T | U | S.obs | SC | Q1 | Q2 | Q3 | Q4 | Q5 | Q6 | Q7 | Q8 | Q9 | Q10 |
| --- | --- | --- | --- | --- | --- | --- | --- | --- | --- | --- | --- | --- | --- | --- |
| Molmol | 71 | 202 | 49 | 0.8823 | 24 | 8 | 3 | 2 | 5 | 0 | 1 | 0 | 1 | 1 |
| Neofur | 21 | 57 | 21 | 0.8488 | 9 | 4 | 4 | 1 | 2 | 0 | 0 | 0 | 0 | 0 |
| Myonig | 44 | 123 | 30 | 0.9135 | 11 | 8 | 2 | 3 | 1 | 0 | 0 | 2 | 1 | 0 |

* Pterygota were excluded from the analyses, reducing the MOTU count by one. As a result, in the following analyses, the diets comprised 49 MOTUs from 71 *M. molossus* (Molmol) samples, 30 MOTUs from 44 *M. nigricans* (Myonig) samples, and 21 MOTUs from 21 *N. furinalis* (Neofur) samples.

* The percentage of sampling completeness for richness (q = 0) is calculated as (1 − Q1 / U) × 100

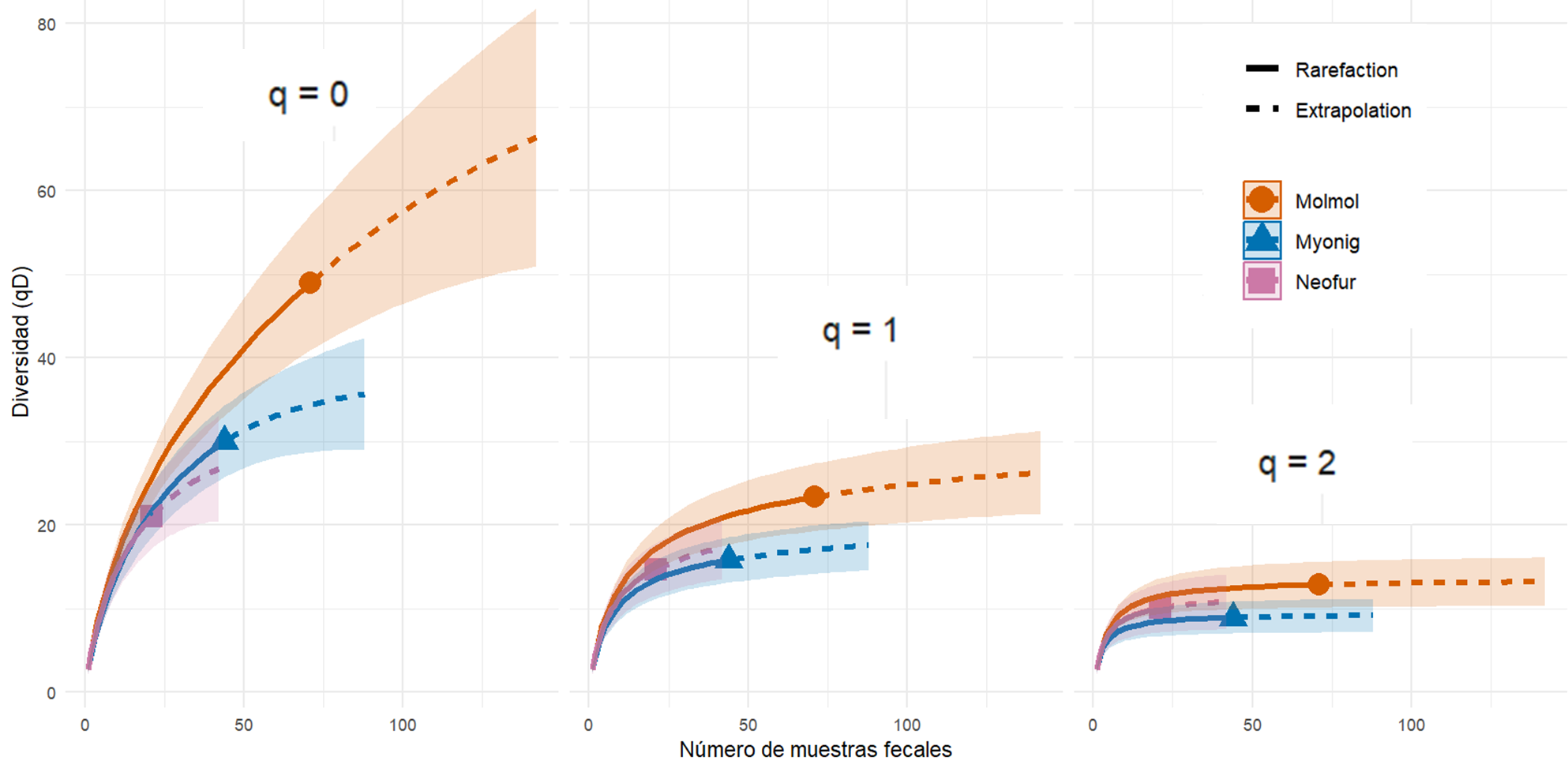

Figure S5a. Rarefaction and extrapolation curves of dietary diversity based on Hill numbers (q = 0, 1, and 2) for *Molossus molossus*, *Myotis nigricans*, and *Neoeptesicus furinalis*.

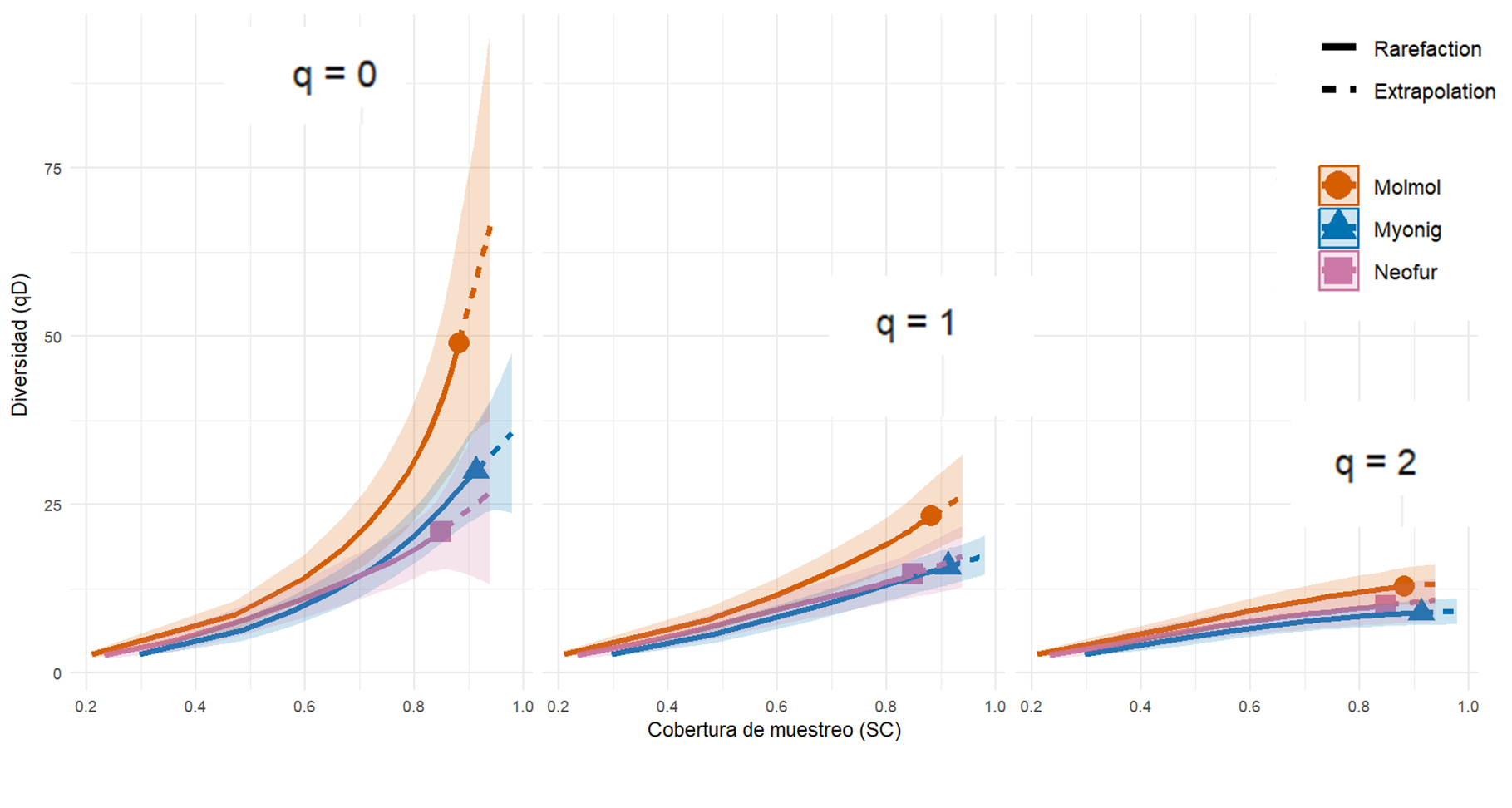

Figure S5b. Coverage‐based rarefaction and extrapolation curves of dietary diversity (Hill numbers q = 0, 1, and 2) for *Molossus molossus*, *Myotis nigricans*, and *Neoeptesicus furinalis*, illustrating sampling completeness (SC) on the x-axis.

**Table SD5c**. Trophic niche breadth of the three bat species based on Hill numbers. Values correspond to taxonomic richness (q₀), Shannon diversity (q₁), and Simpson diversity (q₂) calculated from incidence (presence/absence) dietary data.

| **Especie** | **q₀** | ***q₁** | **q₂** |
| --- | --- | --- | --- |
| Molmol | 49 | 23.3 | 12.9 |
| Neofur | 21 | 14.7 | 10.1 |
| Myonig | 30 | 15.8 | 8.93 |

*q₁ is the main indicator of dietary niche breadth in metabarcoding studies.

**Table S5c**. Pairwise dietary overlap among bat species expressed as local similarity (1 − βₙ) for q₀ (Jaccard-like), q₁ (Horn-like) and q₂ (Morisita–Horn-like).

| **Species 1** | **Species 2** | **q₀** | ***q₁** | **q₂** |
| --- | --- | --- | --- | --- |
| Molmol | Neofur | 0.429 | 0.609 | 0.748 |
| Molmol | Myonig | 0.456 | 0.592 | 0.703 |
| Neofur | Myonig | 0.471 | 0.731 | 0.908 |

*q₁ is the main indicator of dietary niche breadth in metabarcoding studies.
